# Sex differences in intrinsic functional cortical organization reflect differences in network topology rather than cortical morphometry

**DOI:** 10.1101/2023.11.23.568437

**Authors:** Bianca Serio, Meike D. Hettwer, Lisa Wiersch, Giacomo Bignardi, Julia Sacher, Susanne Weis, Simon B. Eickhoff, Sofie L. Valk

**Author notes:** Correspondence to Bianca Serio Sofie L. Valk.

## Abstract

Brain size robustly differs between sexes. However, the consequences of this anatomical dimorphism on sex differences in intrinsic brain function remain unclear. We investigated the extent to which sex differences in intrinsic cortical functional organization may be explained by differences in cortical morphometry, namely brain size, microstructure, and the geodesic distances of connectivity profiles. For this, we computed a low dimensional representation of functional cortical organization, the sensory-association axis, and identified widespread sex differences. Contrary to our expectations, observed sex differences in functional organization were not fundamentally associated with differences in brain size, microstructural organization, or geodesic distances, despite these morphometric properties being *per se* associated with functional organization and differing between sexes. Instead, functional sex differences in the sensory-association axis were associated with differences in functional connectivity profiles and network topology. Collectively, our findings suggest that sex differences in functional cortical organization extend beyond sex differences in cortical morphometry.

**Teaser:** Investigating sex differences in functional cortical organization and their association to differences in cortical morphometry.

## INTRODUCTION

Sex differences in human brain size are robust and widely acknowledged [1–7], but the downstream functional consequences of this anatomical dimorphism are not well understood. Indeed, sex differences in intrinsic brain function are sometimes deemed small or negligible beyond differences attributed to brain size [2]. Nevertheless, diverging patterns of functional connectivity between males and females have been reported even when controlling for differences in brain size and most consistently in sensory and association regions [5, 8, 9]. These regions in fact represent the two anchors of a key principle of hierarchical functional organisation, the sensory-association (S-A) axis, differentiating localized primary sensory/motor areas from a more distributed set of transmodal association regions, including regions belonging to the frontoparietal and default mode networks (DMN) [10, 11]. However, the extent to which sex differences in intrinsic functional cortical organization may be explained by neuroanatomical differences relating to brain size remains unclear.

Brain size and its variability may have important consequences for the spatial distribution of sensory and association areas across the cortical mantle, as illustrated by clear scaling patterns over evolution and development. In fact, over the past 4 million years, hominin evolution has not only shown a general trend of increasing body mass, but also an even more important relative increase in brain size [12]. According to the tethering hypothesis, the brain’s sensory systems, acting as anchors, may have constrained the growth of the developing ancestral mammalian cortex [13]. In this way, evolutionary cortical expansion may have led to the emergence of the S-A axis, with association cortices distributed across the cortical mantle and untethered from sensory hierarchies. Patterns of expansion across cortical regions along the S-A axis are also observed across human development, with a more markedly distributed areal expansion across frontoparietal association regions relative to limbic and sensorimotor areas [14]. Through the increase of overall brain size, the differential expansion of sensory and association areas could thus be an important product of mammalian evolution and development. It is however unclear whether brain expansion and associated re-organization along the S-A axis may also extend to sex differences in cortical morphometry (i.e., cortical shape and size), and thus result in different functional organization of sensory and association regions in males and females.

Morphometric differences between male and female brains have been extensively reported, with males showing a greater absolute brain volume by 8-13% on average [6]. It must be noted that within-group variance in cortical morphometry –which is typically greater in males– is larger than between-group mean effects, meaning that individual differences within sex are larger than group-differences between sexes [15]. Nevertheless, contrary to the belief that males may have larger brains as a sole consequence of their larger bodies [2], it has been repeatedly reported that sex differences in brain size cannot be fully explained by differences in body dimensions, as quantified by height and/or weight [1, 4, 5, 7]. Although individual differences in total brain size seem to account for most differences in relative regional volumes [3], some sex differences still remain statistically significant when the variance explained by total brain size is taken into account [7]. Therefore, there may be sex differences in the scaling of regional brain volume that go beyond linear associations with overall brain and body size. In fact, sex differences in cortical morphometry are partly located at the anchors of the S-A axis [6, 16]. Developmental trajectories of anatomical change also appear regionally heterogeneous, with higher rates of global cortical thickness change found in fronto-temporal association regions and lower rates found in sensory regions [17]. Morphometric cortical properties therefore seem to not only follow patterns of variation along the S-A axis, but also differ between the sexes. Yet, how exactly sex-specific differences in cortical morphometry may be relevant to differences in intrinsic brain function has not been directly explored.

Consistent with patterns of morphometric variation and sex differences, robust evidence points to sex differences in intrinsic functional connectivity (FC) at the poles of the S-A axis [5, 8, 9]. In fact, despite generally controversial findings on sex differences in brain function, findings of stronger FC in females within the DMN [18–20] and in males within sensorimotor areas [19, 21] are consistent and robust. Overlapping morphometric and functional patterns of sex differences along the S-A axis thus suggest that differentiation in functional cortical organization may be somewhat orchestrated by the cortical mantle’s morphometric properties. Indeed, the structure, size, and shape of the cortex not only physically support functional connections, but also determine their length. Short- and long-range connections, as measured by geodesic distance (the distance separating two regions along the cortical mantle) have in fact been found in sensory and association regions respectively [22], thus also displaying patterns of variation along the S-A axis. With increasing distance between regions, cortical function also appears to change more rapidly in association regions relative to sensorimotor regions [23]. These patterns further mirror patterns of microstructural cortical variability identified by post-mortem histology [22] and myelin-sensitive in vivo magnetic resonance imaging (MRI) [22, 23]. As such, intrinsic functional activity, showing variability between the sexes and along the S-A axis, seems to be embedded within the cortical mantle and its microstructural organization. Accumulating evidence further supports the important role played by cortical geometric properties, including size and shape, in sculpting functional architecture. Established findings from graph theory suggest that a cortical functional network’s properties are largely determined by its spatial embedding, namely by the length of its connections [24]. Peaks of DMN clusters on the S-A axis also appear to be equidistantly distributed relative to primary areas [10], in line with the hypothesised untethering of association cortices from sensory hierarchies during evolutionary expansion [13]. Furthermore, recent findings suggest that the spatial organization of intrinsic cortical functional activity is dominated by long wave-lengths of geometric eigenmodes [25]. This research builds on notions from neural field theory positing that brain shape physically constrains brain-wide functional dynamics by imposing boundaries on emerging functional signals [26, 27]. In the context of sex differences in functional cortical organization, brain size also explains some –although not all– sex-specific variance in FC [28]. Together, these findings point to possible morphometric properties that may not only underpin cortical functional architecture, but also be at the root of sex differences in functional organization.

In the current work, we therefore investigated the extent to which sex differences in intrinsic functional cortical organization may be explained by differences in cortical morphometry, namely brain size, microstructure, and the geodesic distances of connectivity profiles. To this end, we used multimodal imaging data (including resting state functional MRI and structural T1 and T2 images) of the Human Connectome Project (HCP) S1200 release [29], consisting of healthy young adults. We began by computing the S-A axis as our measure of functional organization, given its relevance to cortical morphometry and sexual dimorphisms, and tested for sex differences along this low dimensional hierarchical organizational axis. Then, we identified the cortical morphometric properties potentially constraining the S-A axis, including brain size, microstructural organization (a low dimensional microstructure profile covariance (MPC) axis), and the mean geodesic distance of connectivity profiles. Next, we probed associations between patterns of sex differences in cortical morphometry and patterns of sex differences in the S-A axis. Contrary to our expectations, we did not find evidence supporting a morphometric explanation of sex differences in functional organization. As such, we further probed potential functional features that may intrinsically underpin sex differences on the S-A axis, and our findings suggest that differences in FC profiles and network topology may be a more plausible explanation of sex differences in functional organization.

## RESULTS

### Sex differences in the S-A axis of functional cortical organization (Figure 1)

We computed the S-A axis at the individual level as our measure of functional organization in subjects of the HCP S1200 release [29]. For this, we applied a non-linear dimensionality reduction algorithm on functional connectivity (FC) Fisher r-to-z transformed matrices. We only considered the top 10% of the row-wise z-values, representing each seed region’s top 10% of maximally functionally connected regions [30, 31]. We thus found the well-replicated low dimensional axis of functional brain organization explaining the most variance in the data (21.86%) –spanning from unimodal (sensory, here particularly visual) regions to heteromodal (association) regions [10]– and defined it as the S-A axis (Fig. 1A). Then, to test for regional effects of sex on S-A axis loadings (Fig. 1B), we fitted a linear mixed effects model (LMM) including fixed effects of sex, age, and total SA, and random nested effects of family relatedness and sibling status (see Methods for more information on the nested structure of the HCP data and the statistical modelling). We identified sex differences in the S-A axis that were distributed across the seven intrinsic functional Yeo networks [32] (Fig. 1C). Positive *t*-values, depicted in blue, represent higher loadings in males relative to females on the S-A axis, whereas negative *t*-values, depicted in red, represent higher loadings in females relative to males. In Supplementary Figure S1, we also show that patterns of within-sex variability in S-A axis loadings are similar between males and females, with only a few regions showing statistically significant sex differences in variance.

**Figure 1.**
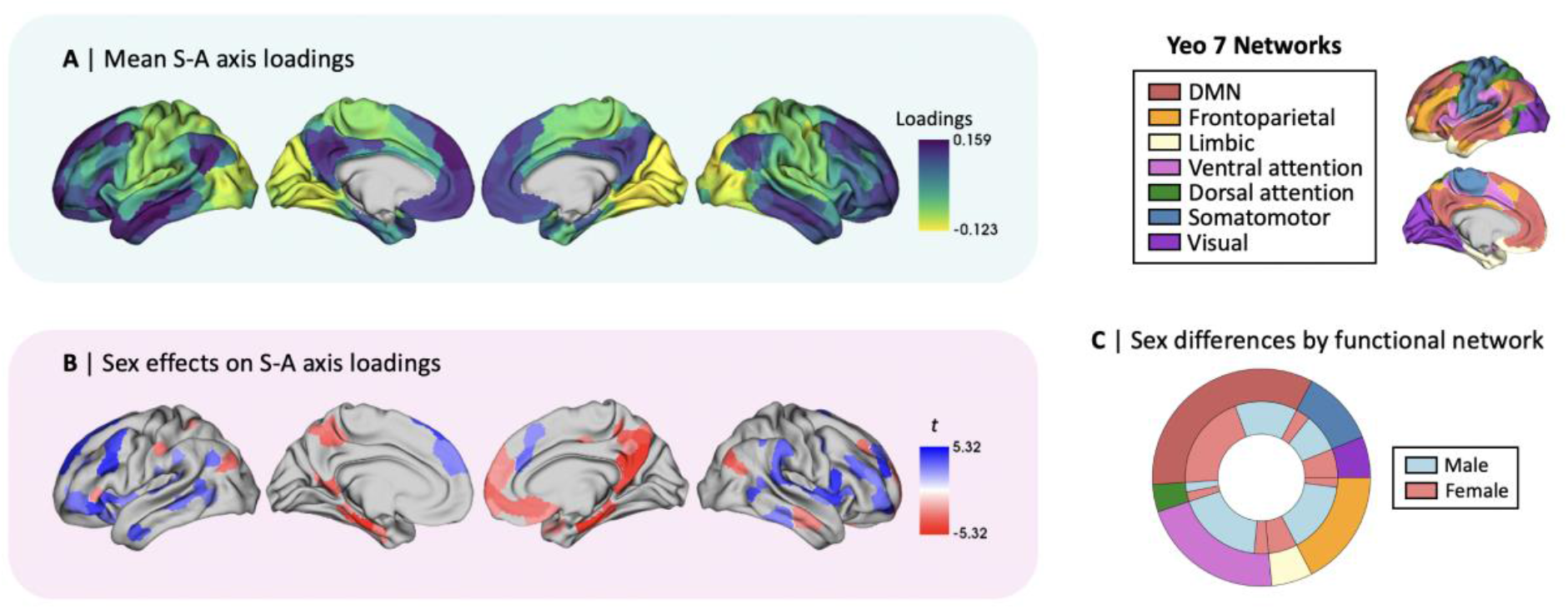
The sensory-association (S-A) axis of functional cortical organization and its sex differences. **A** | Mean S-A axis loadings (spanning from visual to DMN regions) across sexes; **B** | Thresholded *t*-map of linear mixed effect model (LMM) results showing false discovery rate (FDR)-corrected (*q* < 0.05) statistically significant effects of sex on the S-A axis, where blue represents higher male loadings and red represents higher female loadings; **C |** Functional network breakdown of parcels showing statistically significant sex differences in S-A axis loadings. The outer ring displays absolute proportions of statistically significant parcels by functional Yeo network, the inner ring displays absolute proportions by directionality of effects, where blue represents higher male loadings and red represents higher female loadings.

### Morphometric correlates of the S-A axis (Figure 2)

We then investigated potential morphometric constrains of functional organization by probing associations between the S-A axis and brain size, microstructural organization, and the mean geodesic distance of connectivity profiles.

**Figure 2.**
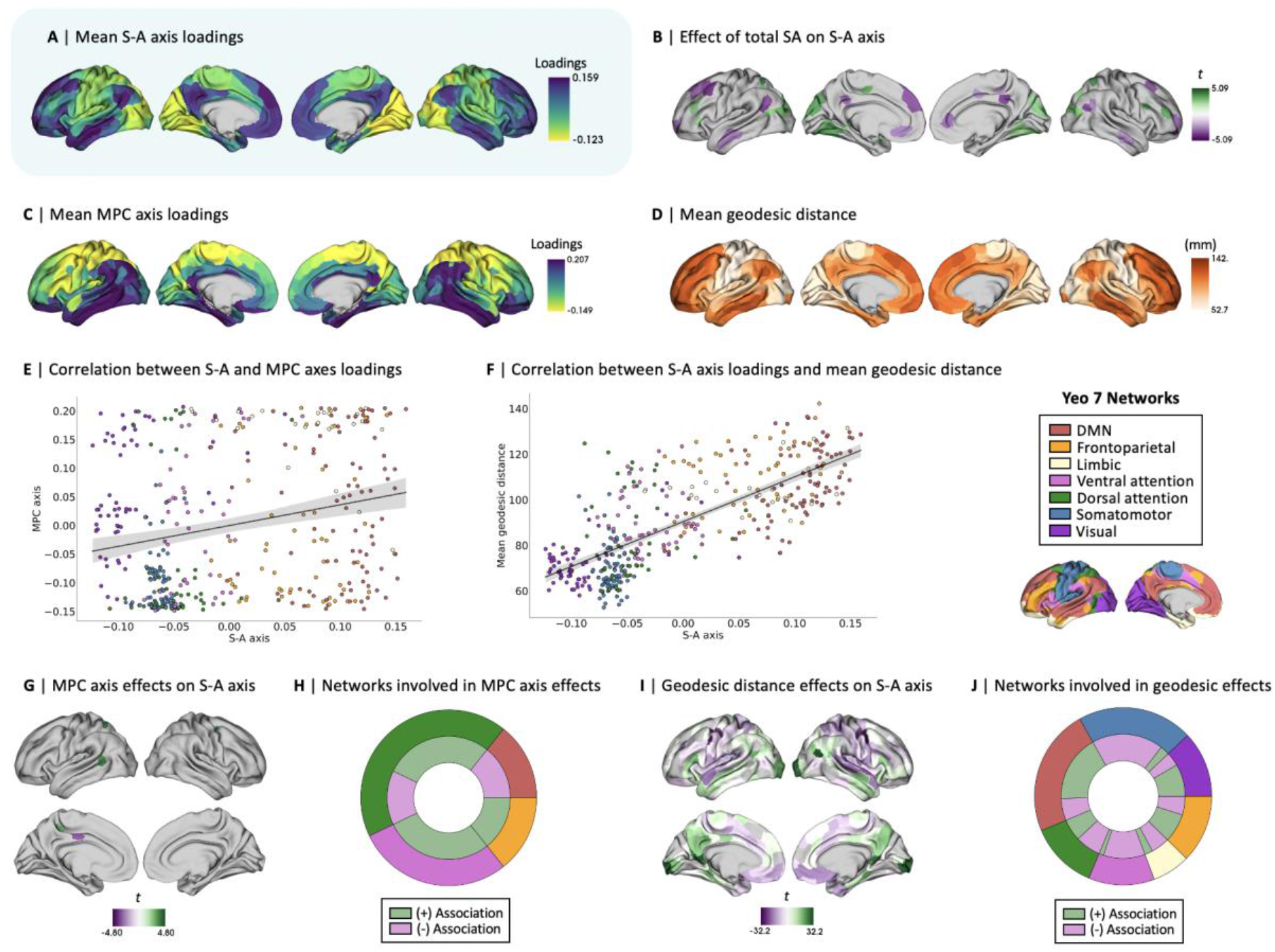
Morphometric correlates of the sensory-association (S-A) axis of functional cortical organization across sexes. **A** | Mean S-A axis loadings (spanning from visual to DMN regions) across sexes; **B** | Statistically significant effects following false discovery rate (FDR) correction (*q* < 0.05) of linear mixed effect model (LMM) results showing total surface area (SA) effects on the S-A axis; **C** | Mean microstructural profile covariance (MPC) axis loadings (spanning from sensory to paralimbic regions) across sexes; **D** | Mean geodesic distance of connectivity profiles across sexes; **E** | Spatial correlation between mean patterns of S-A axis loadings and mean patterns of MPC axis loadings (color-coded by yeo network), *r* = 0.23, *p_spin_* = .024; **F** | Spatial correlation between mean patterns of S-A axis loadings and mean patterns of mean geodesic distance (color-coded by yeo network), *r* = 0.76, *p_spin_* < .001; **G** | Thresholded *t*-map of LMM results showing FDR-corrected statistically significant effects of MPC axis loadings on the S-A axis; ; **H**| Functional network breakdown of parcels showing statistically significant MPC axis effects on S-A axis. The outer ring displays absolute proportions by functional Yeo network, the inner ring displays absolute proportions by directionality of effects; **I** | Thresholded *t*-map of LMM results showing FDR-corrected statistically significant effects of mean geodesic distance on the S-A axis; **J** | Functional network breakdown of parcels showing statistically significant geodesic distance effects on the S-A axis.

First, we tested for associations between the S-A axis loadings and three measures of brain size commonly used in the literature, namely intracranial volume (ICV), total brain volume (TBV), and total surface area (SA). More specifically, ICV represents the entire volume encapsulated by the cranium (i.e., including cerebrospinal fluid), TBV represents the total volume of grey and white matter structures within the neocortex (excluding subcortical structures), and total SA represents the entire SA of the neocortical mantle (see Methods for the exact computation of these measures). Sex differences in brain size and other anthropometric measurements (height, weight, and body mass index) are further reported in Supplementary Table S1. For each measure of brain size, we fitted an LMM to test for regional effects of brain size on S-A axis loadings, and we found total SA to have the most widespread effects amongst the three tested brain size measures (Fig. 2B; Fig. S2).

Second, we computed a MPC axis of organization at the individual level, representing a low dimensional representation of the similarity of T1-weighted (T1w) over T2-weighted (T2w) tissue intensity across cortical regions and layers [33–35]. We computed the MPC axis by again conducting non-linear dimensionality reduction on MPC matrices [30, 31], which were obtained by sampling and correlating the intracortical microstructural intensity of 12 equivolumetric depth profiles (see Methods). Following the same approach used for computing the S-A axis, we selected the axis explaining the most variance in the data (25.97%) –spanning from sensory to paralimbic regions– defining it as the MPC axis (Fig. 2C). We specifically selected this low-dimensional representation of microstructural organization as it has been previously shown to covary with the low-dimensional representation of functional organization (i.e., the S-A axis) [34]. To test for whole-brain associations between the S-A and MPC axes, we correlated the spatial maps of the axes’ mean loadings (Fig. 2A and 2C) across all subjects (Fig. 2E; *r* = 0.23, *p_spin_* = .024). We further fitted an LMM to test for regional effects of MPC axis loadings on S-A axis loadings at the parcel level (Fig. 2G and 2H), and found small and localized associations between the S-A and MPC axes.

Third, we computed the mean geodesic distance of connectivity profiles at the individual level. The mean geodesic distance of connectivity profiles is the mean distance along the cortical mantle between each region and its top 10% maximally functionally connected regions. Group-level patterns (i.e., averaged across all subjects; Fig. 2D) revealed shorter distances in visual and somatomotor (sensory) regions, and longer distances in frontoparietal and DMN (association) regions. We also tested for whole-brain associations between the S-A axis and patterns of mean geodesic distance of connectivity profiles by correlating their spatial maps (Fig. 2A and 2D) averaged across all subjects (Fig. 2F; *r* = 0.76, *p*_spin_ < .001). We again also fitted an LMM to test for regional effects of mean geodesic distance on S-A axis loadings at the parcel level (Fig. 2I and 2J) and found strong and widespread associations between the S-A axis and patterns of mean geodesic distance.

### Morphometric correlates do not explain sex differences in the S-A axis (Figure 3)

After establishing the morphometric correlates of the S-A axis, we probed whether sex differences in cortical morphometry may explain sex differences in the S-A axis. First, we tested whether sex differences in S-A axis loadings were moderated by total SA. For this, we modeled an interaction term of sex by total SA on the S-A axis loadings within the original LMM (Fig. 3B) and found no statistically significant effects across regions. In Supplementary Figure S3A-C, we further show that this interaction effect when including height as a covariate to the LMM yields virtually the same *t*-values as when height is not included as a covariate (*r* = 0.99, *p*_spin_ < .001). This suggests that height –being an anthropometric feature that systematically differs between the sexes– does not explain variance in the moderation of sex effects by total SA on the S-A axis loadings either. We also plotted within-sex effects of total SA on S-A axis loadings, showing similar although slightly diverging patterns of effects between males and females (*r* = 0.65, *p*_spin_ = .001; Fig. S3D-F). However, the divergence of patterns between sexes may not be strong or systematic enough to be interpreted as meaningful, as underlined by the lack of evidence of a statistically significant sex by total SA interaction effect on the S-A axis. Second, we tested for regional sex effects on the MPC axis loadings (Fig. 3C) and correlated this spatial *t*-map with the *t*-map depicting regional sex effects on the S-A axis loadings (Fig. 3A). Here, we found no statistically significant association between these two patterns of sex differences (Fig. 3E; *r* = 0.03, *p*_spin_ = .388). Third, we tested for regional sex effects on the mean geodesic distance of connectivity profiles (Fig. 3D) and again correlated this spatial *t*-map with the *t*-map depicting sex effects in the S-A axis loadings (Fig. 3A). Again, we found no statistically significant association between these two patterns of sex differences (Fig. 3F; *r* = 0-.04, *p*_spin_ = .395). These results together suggest that sex differences in the S-A axis are overall not fundamentally moderated by –or associated with– sex differences in cortical morphometry.

**Figure 3.**
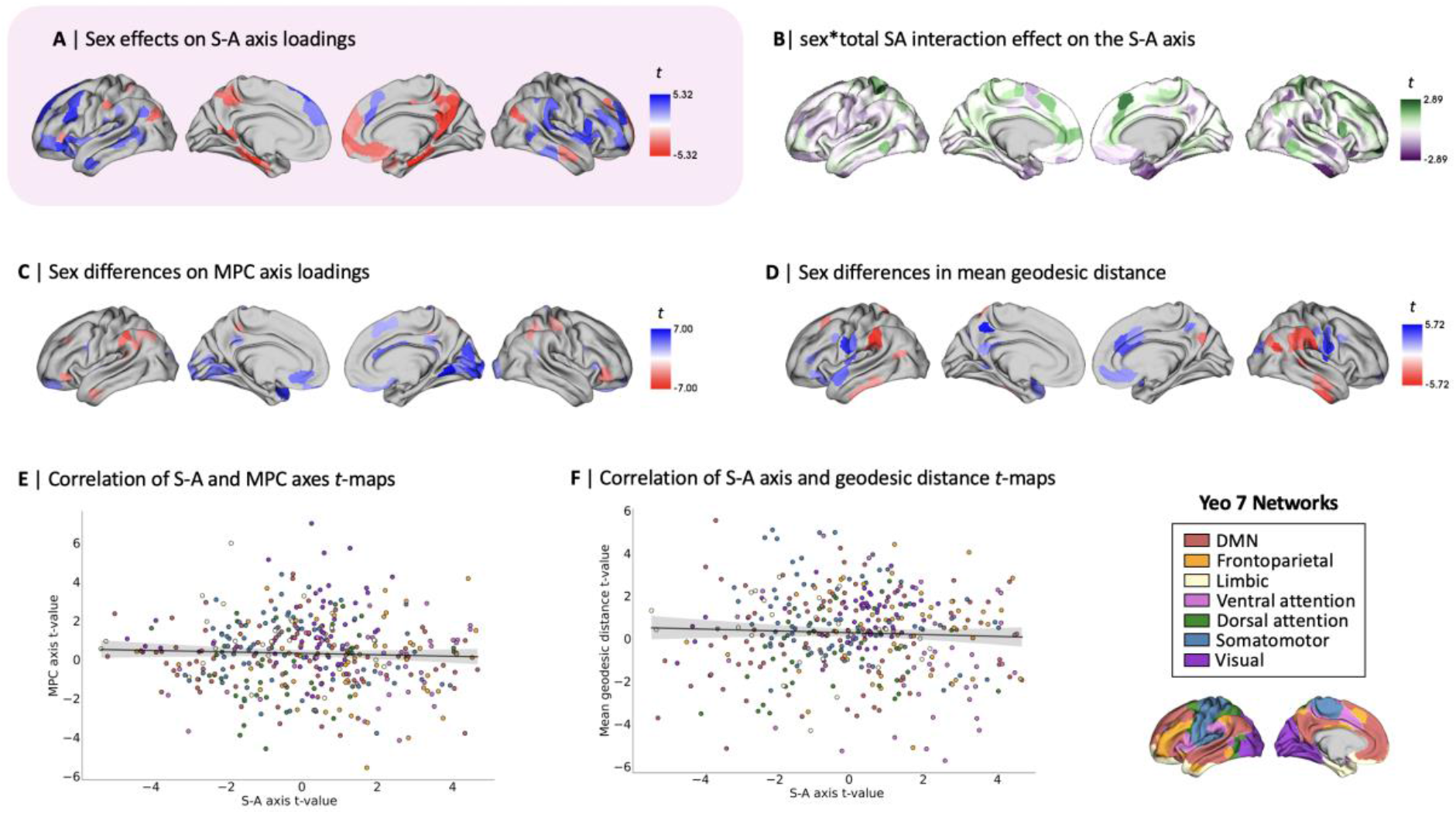
Morphometric correlates of sex differences in the sensory-association (S-A) axis. **A** | Thresholded *t*-map of linear mixed effect model (LMM) results showing false discovery rate (FDR)-corrected (*q* < 0.05) statistically significant effects of sex on the S-A axis, where blue represents higher male loadings and red represents higher female loadings; **B** | Unthresholded *t*-map of LMM testing for sex by total surface area (SA) interaction effects on S-A axis (there were no statistically significant sex effects after FDR correction; **C** | Thresholded *t*-map of LMM results showing FDR-corrected statistically significant effects of sex on the microstructure profile covariance (MPC) axis; **D** | Thresholded *t*-map of LMM results showing FDR-corrected statistically significant effects of sex on the mean geodesic distance of connectivity profiles; **E** | Scatterplot displaying the spatial correlation between patterns of sex differences (*t*-maps) in S-A axis loadings and in MPC axis loadings (color-coded by yeo network), *r* = 0.03, *p_spin_* = .388; **F** | Scatterplot displaying the spatial correlation between patterns of sex differences (*t*-maps) in S-A axis loadings and in the mean geodesic distance of connectivity profiles (color-coded by yeo network), *r* = -0.04, *p_spin_* = .395.

As an additional sensitivity analysis, we found that including the MPC axis and the mean geodesic distances as covariates in our LMM testing for sex effects on the S-A axis yields highly similar regional sex effects to those reported in Figure 1A (for which the original LMM only included total SA as a morphometric covariate, to control for brain size), as shown by the strong correlation of *t*-maps (*r* = 0.95, *p_spin_* < .001, Supplementary Fig. S4A). Similarly, the association between sex effects when including all morphometric covariates versus not including any (i.e., also excluding total SA) remains high despite a small decrease in correlation strength (*r* = 0.81, *p_spin_* < .001, Supplementary Fig S4B). These findings further suggest that sex differences in brain size only explain a minor amount of variance in sex differences in the S-A axis.

### Intrinsic functional underpinnings of differences in the S-A axis (Figure 4)

Given that sex differences in the morphometric correlates of the S-A axis did not appear to explain sex differences in the S-A axis, we probed potential intrinsic functional underpinnings of sex differences on the S-A axis. We thus tested for associations between sex differences in the S-A axis loadings and sex differences in mean FC strength, FC profiles, and network topology.

First, we computed mean FC strength at the individual level from FC matrices, representing –for each parcel– the mean row-wise z-values of a given seed region’s top 10% maximally functionally connected regions. We then fitted an LMM to test for local effects at the parcel level of sex on mean FC strength (Fig. 4C and 4E), which revealed –amongst other sex differences– higher intrinsic FC in females in DMN regions and in males in somatomotor regions. To test associations between patterns of sex differences in the S-A axis loadings and in FC strength, we spatially correlated the *t*-maps (Fig. 4A and 4C) of the respective sex differences and did not detect a statistically significant association between sex differences in the S-A axis and sex differences in FC strength (Fig. 4H; *r* = 0.02, *p*_spin_ = .380).

**Figure 4.**
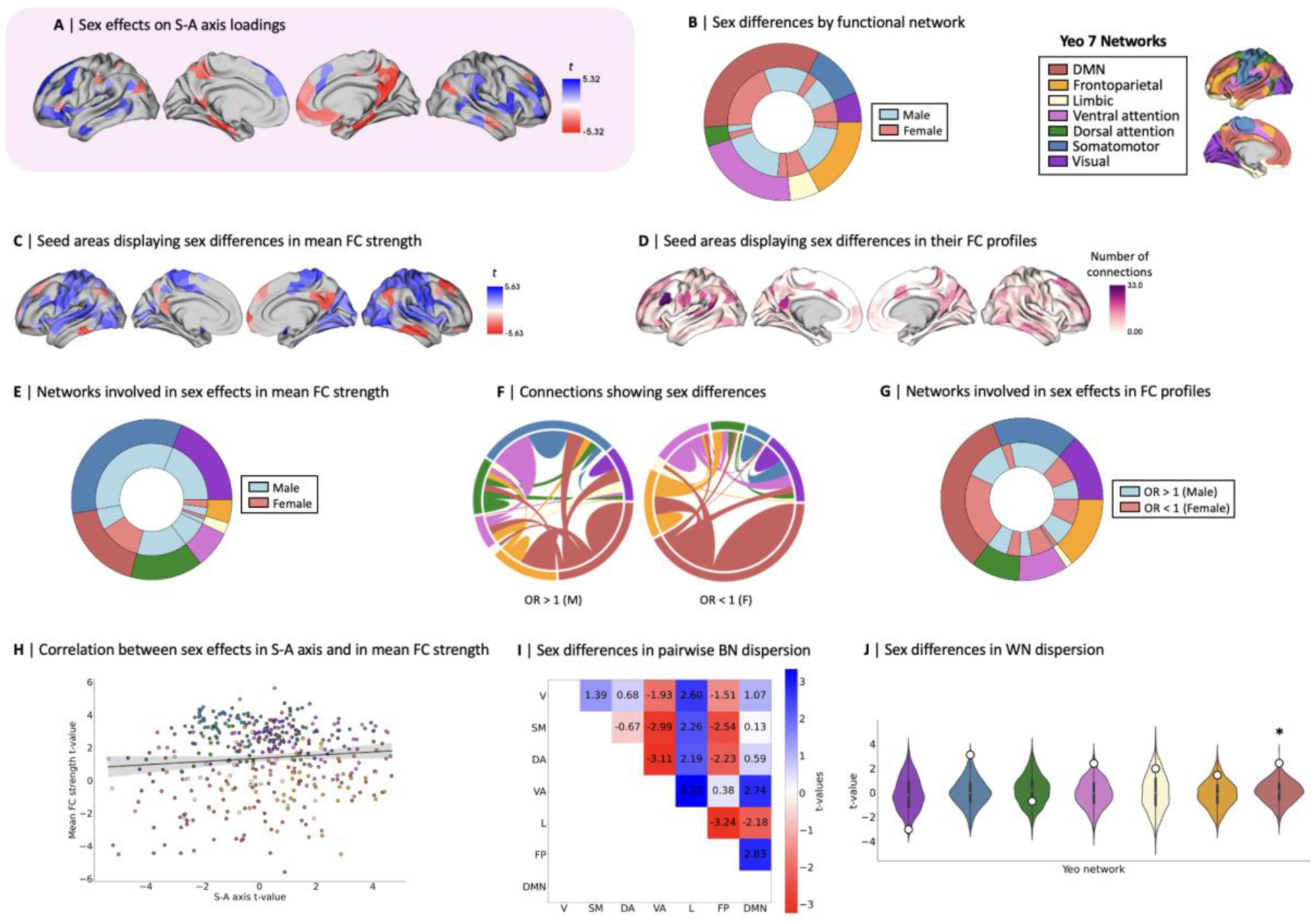
Intrinsic functional underpinnings of differences in the sensory-association (S-A) axis. **A** | Thresholded *t*-map of linear mixed effect model (LMM) results showing false discovery rate (FDR)-corrected (*q* < 0.05) statistically significant effects of sex on the S-A axis, where blue represents higher male loadings and red represents higher female loadings; **B** | Functional network breakdown of parcels showing statistically significant sex differences in S-A axis loadings. The outer ring displays absolute proportions of statistically significant parcels by functional Yeo network, the inner ring displays absolute proportions by directionality of effects, where blue represents higher male loadings and red represents higher female loadings; **C** | Thresholded *t*-map of LMM results showing FDR-corrected statistically significant effects of sex on mean functional connectivity (FC) strength; **D** | Number of connections (per seed region) showing statistically significant FDR-corrected sex differences in their odds of belonging to the given seed’s top 10% of connections; **E** | Functional network breakdown of connections showing statistically significant FDR-corrected sex differences in mean FC strength; **F** | Connections between seed and target regions showing statistically significant FDR-corrected sex differences in FC profiles (OR > 1 meaning that males have higher odds than females of having a target region belong to a seed region’s top 10% connections, where OR < 1 means that females have higher odds than males of having a target region belong to a seed region’s top 10% connections; connections are color coded by network and weighed by number of connections between the network pairs; **G** | Functional network breakdown of connections showing statistically significant FDR-corrected sex effects in their odds of belonging to the given seed’s top 10% of connections; **H** | Spatial correlation between patterns of sex effects in S-A axis loadings and patterns of sex effects in mean FC strength (color-coded by yeo network), *r* = 0.02, *p*_spin_ = .380; **I** | *t*-values for the sex contrast in between-network (BN) dispersion for each pairwise Yeo network comparison, where blue represents higher male BN dispersion and red represents higher female BN dfispersion (no statistically significant sex effects after spin permutation and Bonferroni correction; *p*_spin_ < .001); **J** | *t*-values for the sex contrast in within-network (WN) dispersion for each Yeo network (displayed as white dots), plotted on null distributions of *t* statistics derived from 1000 spin permutations, where positive *t*-values represent higher male WN dispersion and negative *t*-values represents higher female WN dispersion, * indicates Bonferroni-corrected (*p*_spin_ < .004) statistical significance of the sex contrast. V, visual, SM, somatomotor, DA, dorsal attention, VA, ventral attention, L, limbic, FP, frontoparietal, DMN, default mode network.

Second, we defined FC profiles at the individual level, for which we identified the top 10% of maximally functionally connected regions. Using the Chi-square (χ^2^) test of independence, we assessed –for each possible pairwise connection along the 400x400 matrix– sex differences in a given target region’s odds of belonging to the top 10% of maximally functionally connected regions of a given seed region. We identified the direction of these sex effects with the odds ratio (OR), where OR > 1 indicates a given region’s greater male odds and OR < 1 indicates a given region’s greater female odds. Out of the 160000 tested functional connections, 2004 connections (corresponding to 1.25% of all connections) displayed statistically significant sex differences in their odds of constituting a seed’s top 10% connections after FDR correction (Fig. 4F), suggesting that sex differences in S-A axis loadings may in part stem from differences in FC profiles, namely differences in which functional connections are the strongest. For illustrative purposes, we summarized spatial patterns of sex differences in FC profiles as the sum of connections showing sex differences per seed region (Fig. 4D), as well as the overall networks involved in sex differences in FC profiles (Fig. 4G).

Finally, we investigated network topology, namely the organization of networks along the S-A axis. We computed between-network dispersion for each subject, quantifying the pairwise distance between two given networks along the S-A axis, where a higher value indicates higher segregation of the given pair of networks (21 pairs of Yeo networks in total) [36]. We also computed within-network dispersion for each subject for the seven intrinsic networks under study [32], quantifying the spread of regions within each network along the S-A axis, where a higher value indicates higher segregation of the given network’s regions. LMMs did not show any statistically significant sex differences in between-network dispersion for any of the network pairs (Fig. 4I). However, we found greater male within-network dispersion in the DMN, *t* = 2.41, *p*_spin_ = 0.001 (Fig. 4J), revealing a greater spread of regions belonging to the DMN along the S-A axis in males. The full statistical results for the analysis of sex differences in network dispersion are summarized in Supplementary Table S2.

## DISCUSSION

In the current work, we investigated the extent to which sex differences in functional cortical organization may be explained by differences in cortical morphometry, namely brain size, microstructure, and the geodesic distances of connectivity profiles. We identified widespread sex differences in adult functional cortical organization as defined by the S-A axis, which however did not appear fundamentally associated with sex differences in brain size, microstructural organization, nor the mean geodesic distance of connectivity profiles. This finding is particularly striking given that the morphometric properties under study were all *per se* associated with the S-A axis and differed between sexes. Instead, we observed that sex differences in the S-A axis were related to differences in FC profiles and network topology, namely greater male dispersion within the DMN. Collectively, our findings suggest that sex differences in functional cortical organization go beyond neuroanatomical sex differences pertaining to cortical morphometry.

Different measures of brain size are commonly used in the literature, including ICV, TBV, and total SA. Although these measures highly covary and are often used interchangeably, they quantify different morphometric features of the brain, with sex differences in “brain size” ranging from 8% to 13% depending on the selected measure [6]. The size and direction of sex effects also vary by neuroanatomical property, such as different tissue types, brain regions, and features (including cortical thickness, gyrification, and SA) [37]. Furthermore, morphometric features vary differently as a function of age, whereby for example TBV but not ICV is affected by atrophy [6]. These findings highlight the complex heterogeneity of neuroanatomical properties constituting brain size. The potential for introducing non-linear bias in the detection of sex effects should therefore not be overlooked, particularly when statistically controlling for brain size in the detection of sex effects on brain structure and function [28, 38–41]. We addressed this issue by testing the effects of different measures of brain size, namely ICV, TBV, and total SA, on the S-A axis. Here, given that total SA had the most widespread effects on functional organization, we deemed it the most appropriate measure of brain size, which we further included as a covariate in our models throughout our analyses. The relevance of total SA is also supported by the theoretical assumptions motivating our study, namely the relevance of cortical shape and geometry in constraining brain wide functional dynamics [25–27] and thus sex differences in these features potentially underpinning sex differences in the S-A axis. Our findings therefore highlight the diverging effects of different measures of brain size and depict total SA as having the most substantial theoretical and statistical associations to a low dimensional representation of functional cortical organization. As such, future research on sex differences should also carefully select the measure of brain size that is most conceptually and empirically pertinent to the research question under study in order to minimize bias.

By establishing morphometric correlates of the S-A axis in addition to brain size, namely a microstructural axis of cortical organization [34, 35, 42] and the mean geodesic distance of connectivity profiles [10, 42], our findings align with previous work and argue for the rooting of functional cortical organization in cortical structure and shape. We show a particularly strong association between the mean geodesic distance of connectivity profiles and the S-A axis, supporting the relevance of the cortical mantle’s shape in sculpting functional organization. This may be a product of the cortical mantle’s evolutionary expansion, where association regions are untethered from sensory hierarchies [13], and long-range connections preserve the overall connectedness of cortical networks by facilitating the communication between distant areas [24]. Furthermore, as indexed by the MPC axis, microstructural organization appears to mildly covary with the S-A axis, supporting to some degree the well-established idea of structural constraints on brain function [34, 35, 43]. In our study, we obtained intensity profiles via the ratio of T1w over T2w imaging sequences, and although it is commonly used to measure myelin [34, 35, 44], the T1w/T2w ratio has been described as an acceptable *qualitative* proxy for myelin in grey but not white matter [45]. It is indeed thought to capture unique features of microstructural tissue that appear largely independent of diffusion-based metrics, thus portraying a mix of neuroanatomical features beyond pure myelin [46]. We therefore consider the T1w/T2w ratio –and the resulting MPC axis– as a general measure of tissue microstructure, which may serve as a scaffold for functional organization.

After establishing morphometric correlates of the S-A axis, we addressed our primary aim of probing the extent to which sex differences in functional cortical organization may be explained by sex differences in cortical morphometry. We observed slightly diverging results when including – as opposed to excluding– total SA as a covariate in our model testing for sex differences in S-A axis loadings, suggesting that sex differences in total SA explained some variance in sex differences in functional organization. This finding is consistent with the systematic practice of controlling for brain size when testing for structural and functional sex differences [28, 38–41]. Nevertheless, our findings overall suggest that morphometric differences between the sexes are altogether not substantial contributors of sex differences in the S-A axis of functional organization. We did not find sex differences in S-A axis loadings to be moderated by total SA, nor any associations between patterns of sex differences in the S-A axis and patterns of sex differences in the MPC axis or in the mean geodesic distance of connectivity profiles. The negligeable relevance of cortical morphometry to sex differences in the S-A axis is striking given that morphometric properties appear *per se* to be associated with the S-A axis and to differ between sexes. The mechanisms underpinning different patterns of morphometric and functional sex differences may thus be independent from one another, suggesting that sex differences in functional cortical organization may extend beyond the connectome’s supporting shape and structure.

Given that sex differences in morphometric correlates of the S-A axis did not seem to explain sex differences on the S-A axis, we probed and found potential intrinsic functional underpinnings of sex differences on the S-A axis. Firstly, the sex differences we observed in the S-A axis loadings were distributed across functional networks, and notably in the DMN, frontoparietal and ventral attention networks. This is consistent with previous findings of greater individual variability in the functional topography of these association networks relative to lower-order sensory networks, which have also been shown to contribute the most to sex classification in youth [8]. We also observed sex differences in intrinsic FC strength, replicating previous widely established findings of greater FC in females within DMN regions [18–20] and in males within somatomotor regions [19, 21]. However, these patterns did not spatially overlap with patterns of sex differences in the S-A axis, suggesting that FC strength is not a feature of intrinsic FC that is captured by sex differences in our low dimensional representation of functional organization. Instead, we found that sex differences in the S-A axis were related to differences in FC profiles, which also presented qualitative sex differences in the proportional breakdown of networks involved. Females seemed to make more top connections involving the DMN relative to males, whereas males displayed more top connections involving the somatomotor networks relative to females. These sex differences in the configuration of connections may not only underly the recurrence of sex differences in these networks [18–21], but may also explain sex differences in network topology.

We in fact observed greater male dispersion (i.e., decreased similarity on the S-A axis) within the DMN (and the somatomotor network barely not surviving the Bonferroni correction), which is also consistent with previous findings of generally more segregated male networks [47]. These network-specific topological sex differences may be related to greater female odds of connections within the DMN, and greater male odds of somatomotor connections with other networks. Concretely, network topology, which represents the organization of functional communities within and between functional networks [36], may reflect brain states [48]. Network topology has also been associated with different cognitive features including arousal [49], awareness and consciousness [50], behavior and task performance [51], and cognitive flexibility [52]. The balance between integration and segregation is complex, dynamic, and necessary to maintain the brain’s metastability [53] by reaching a point of equilibrium between global organization and local specialization [43]. The brain is a highly interconnected and metabolically expensive organ, and its organization is required to dynamically balance topological efficiency and energy utilization in response to transient cognitive and physiological demands [54]. Our findings of sex differences in network topology may therefore pertain to intricate sex differences not only in brain states at rest, which may underpin cognitive differences, but also in energy expenditure, which would reflect physiological differences.

Despite the novel insights gained through our study, some limitations must be acknowledged. Firstly, by only considering biological sex, we neglected possible effects of gender on functional organization and its morphometric correlates. Findings may indeed appear more nuanced if we moved beyond the unrealistic assumption of a clear-cut sexual dimorphism of brain structure and function [55], as the relevance of considering transgender individuals in the study of sex differences is being increasingly recognized [56]. Nevertheless, we intentionally focused on the biological and dichotomous variable of sex assigned at birth given that our study aimed to study biological mechanisms relating to cortical morphometry. We did not venture in the intricacies of gender as they require an additional careful consideration of complex social and environmental influences, which go beyond the scope of our study. Secondly, we focused on neocortical functional organization, excluding subcortical structures and the cerebellum despite their substantial contributions to whole brain organization through their notable structural integration with the cortex [57]. In fact, the amygdala and hippocampus are hypothesized to be at the origin of mammalian cortical evolution [58] and have also repeatedly shown both structural [6] and functional [59] sex differences. Nevertheless, our exclusive focus on the neocortex was motivated by the relevance of using the S-A axis as our measure of functional organization, which is obtained by reducing the dimensionality of FC matrices of cortical data [10]. By using the S-A axis, our work identified sex differences embedded in a key macroscale organizational principle that is closely tied to evolutionary expansion and cortical morphometry, going beyond previous research on functional differences between the sexes that solely focus on intrinsic brain function. Thirdly, the morphometric properties considered in our study are not exhaustive, overlooking the contributions of other morphometric measures such as local volumes of grey matter. The inclusion of the MPC axis [34, 35] and the mean geodesic distance of connectivity profiles [10, 42] was however supported by their theoretical and empirical relevance to functional cortical organization, particularly its low dimensional embedding.

All in all, our study opens a new set of questions pertaining to the mechanisms underpinning sex differences in functional cortical organization, given that they do not appear to be fundamentally rooted in cortical morphometric differences. Our findings instead suggest that sex differences in the S-A axis are to some extent intrinsically related to differences in FC profiles and network topology. Therefore, future research should explore factors driving males and females to form distinct functional connections and to adopt divergent system-level organization of functional networks. Recognizing the human body as a complex system of systems, future work should investigate other biological factors that may contribute to functional sex differences such as genes located on sex chromosomes [16] and sex hormones [60, 61]. Environmental factors should equally be considered, for example stress, which has also been found to contribute to sex differences via epigenetic mechanisms [62]. Investigating the mechanisms underpinning sex differences in functional organization is crucial to gain a deeper understanding of discrepancies in predisposition to psychiatric disorders, for example greater female vulnerability to affective disorders throughout the lifespan [63, 64] and particularly during hormone transition periods such as puberty, pregnancy, and menopause [65].

## MATERIALS AND METHODS

### Participants and Experimental Design

Our analyses were conducted on the publicly available data of healthy young adults from the Human Connectome Project (HCP) S1200 release (http://www.humanconnectome.org/) [29]. We selected subjects with available functional, T1, and T2 data, resulting in a final sample of 1000 individuals (536 females) with a mean age of 28.73 ± 3.71 years, including 284 monozygotic twins (MZ), 184 dizygotic twins (DZ), 443 non-twin siblings, and 89 unrelated individuals. Subjects were all born in Missouri but recruited in an attempt to broadly reflect the racial and ethnic composition of the United States population. Recruitment efforts aimed to yield a subject pool capturing a wide range of variability –in socioeconomic and behavioral terms– in order to be representative of the general healthy population. The term “healthy” was thus broadly defined. Individuals with documented neurodevelopmental and psychiatric disorders, or reporting physiological illnesses such as high blood pressure or diabetes were excluded, but not individuals who reported smoking, being overweight, or a history of recreational drug use or heavy drinking (if they had not experienced severe symptoms). Informed consent was obtained for all study subjects. More detailed information about the HCP study design and recruitment procedure is available elsewhere [29, 66].

### Structural MRI acquisition and preprocessing

The HCP’s MRI data was acquired on a customized 3T Siemens Skyra ConnectomeScanner with a 32-channel head coil at Washington University across four scanning sessions held over two days. Structural MRI images were acquired on the same day via high resolution T1-weighted (T1w) and T2-weighted (T2w) sequences. Two separate T1w images were acquired and averaged, with identical scanning parameters using a 3D MPRAGE sequence (0.7 mm isovoxels, FOV = 224 mm, matrix = 320 × 320 mm, 256 sagittal slices; TR = 2400 ms, TE = 2.14 ms, TI = 1000 ms, flip angle = 8°, BW = 210 Hz per pixel, ES = 7.6 ms). Two separate T2w images were acquired and averaged, with identical scanning parameters using a variable flip angle turbo spin-echo (3D T2-SPACE) sequence, with the same isotropic resolution, matrix, FOV, and slices as for the T1w sequence (TR = 3200 ms, TE = 565 ms, BW = 744 Hz per pixel, total turbo factor = 314). The preprocessing steps included co-registering the T1w and T2w images, bias field (B1) correction, registration to MNI space, segmentation, and surface reconstruction. See [29, 66, 67] for more detail on the HCP’s MRI protocols and the FreeSurfer segmentation pipeline.

### Functional MRI (fMRI) acquisition and preprocessing

The HCP’s fMRI data was collected after the structural sequences and following the HCP’s minimal processing pipeline, as described above. A total of 1h of resting-state functional data was collected across four identical 15min scanning sessions, equally split over two days (LR1, RL1, LR2, RL2), with a gradient echo EPI sequence at a resolution of 2 mm isotropic (FOV = 208 × 180 mm, matrix = 104 × 90 mm, 72 slices covering the whole brain, TR = 720 ms, TE = 33 ms, multiband factor of 8, FA = 52°). The multimodal surface matching algorithm (MSMAll) was used to co-register the data to the HCP template 32 k_LR surface space, consisting of 32492 nodes per hemisphere (59412 nodes excluding the medial wall). A more detailed description of the resting state fMRI data acquisition and analysis protocol is available elsewhere [67, 68].

### Functional connectivity (FC) and the sensory-association (S-A) axis of functional organization

Throughout this work, we used the Schaefer 400 parcellation (clustered into 7 networks: visual, somatomotor, dorsal attention, ventral attention, limbic, frontoparietal, DMN [32]). This widely used functionally-derived parcellation scheme was originally obtained via a gradient-weighted Markov Random Field model integrating local gradient and global similarity approaches [69]. The vertex-wise functional timeseries were therefore averaged within the Schaefer 400 cortical parcels. FC matrices (400x400) were then computed at the individual level –per scanning session– by correlating cortical timeseries in a pairwise manner using the Pearson product moment. We normalized the correlation coefficients using Fisher’s z-transformation. Final FC matrices were obtained by averaging each subject’s matrices across their four scanning sessions. From these FC matrices and for each subject, we computed the S-A axis of functional organization, as described below.

We conducted data reduction on the FC matrices to yield macroscale gradients of functional organization [10]. For this, we used diffusion map embedding, a non-linear manifold learning algorithm that reduces complex, high-dimensional structures of data (in our case affinity matrices) to low-dimensional representations combining geometry with the probability distribution of data points [30]. Thus, cortical parcels that are strongly interconnected are represented closer together in the resulting low dimensional manifold of FC data, whereas parcels with low covariance are represented farther apart, as indexed by the parcels’ gradient loadings. To this end, we used the BrainSpace Python toolbox [31] to compute 10 gradients with the following parameters: 90% threshold (i.e., only considering the top 10% row-wise z-values of FC matrices, representing each seed region’s top 10% of maximally functionally connected regions), α = 0.5 (α controls whether the geometry of the set is reflected in the low-dimensional embedding – i.e., the influence of the sampling points density on the manifold, where α = 0 (maximal influence) and α = 1 (no influence)), and t = 0 (t controls the scale of eigenvalues). These parameters were selected for consistency with previous studies [10, 34] and represent choices that are recommended to retain global relations between datapoints in the embedded space whilst being relatively robust to noise. In order to increase comparability for further between-subject analyses, Procrustes alignment was used to align individual gradients to mean gradients, which were computed by applying diffusion map embedding –with the same parameters listed above– to the mean FC matrix (i.e., FC matrices averaged across all subjects). The computation of these FC gradients was carried out independently per hemisphere (i.e., considering the top 10% row-wise z-values of only half of the FC matrices, shaped 200x200) and the gradient loadings resulting from both hemispheres were subsequently concatenated. This decision was made for consistency and comparability reasons within our study, so that the top 10% functional connections selected for data reduction corresponded to those considered in the calculation of the mean geodesic distance of connectivity profiles –which were only computed per hemisphere– as described further below). We verified and confirmed the stability FC gradients when computing them per hemisphere versus at the whole brain level, as shown by the spatial correlation of mean gradient loadings (*r* = 0.98, *p*_spin_ = .001). Finally, we took the well-replicated principal gradient explaining the most variance in the data and spanning from visual to DMN regions [10], which we labeled the S-A axis and used to represent functional organization for subsequent analyses.

We also computed, for each subject, mean FC strength at the parcel level in a seed-wise fashion, by averaging the row-wise z-values of each seed region’s top 10% maximally functionally connected regions –again per hemisphere– and subsequently concatenated the hemispheric mean FC strength values to reconstruct whole brain data.

### Cortical microstructure and microstructural profile covariance (MPC)

Microstructural properties –including myelin and cellular characteristics– show depth-dependent variation along cortical columns, as reported by histology [34, 70, 71] as well as *in vivo* and *post mortem* neuroimaging [33–35, 71], which illustrates cortical hierarchy [11]. Similar to previous work [35], we quantified cortical microstructure, or “microstructural profile intensity” (MPI), using the myelin-sensitive MRI contrast obtained from the T1w/T2w ratio from the HCP minimal processing pipeline described above [67] (a reliability check is reported in the Supplementary Methods, Fig. S5). The T1w/T2w ratio uses the T2w image to correct for inhomogeneities in the T1w image [44]. Then, we followed the previously described protocol [33–35] to compute our measurement of MPC, which reflects the variation of MPI, across cortical depths. In short, we generated 14 equivolumetric surfaces within the inner and outer cortical surfaces, then excluded the inner- and outer-most surfaces, thus remaining with 12 surfaces representing cortical layers. Surface generation was based on a model compensating for cortical folding by altering the pairwise Euclidean distance (ρ) of intracortical surfaces throughout the cortex and thus preserving fractional volume between the surfaces. For each surface, ρ was calculated as defined in Eq. 1.

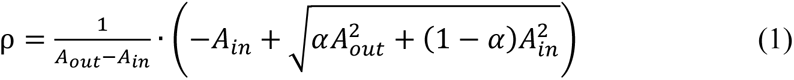

for which *α* denotes a fraction of the total volume of the segment that the surface accounts for, while *A*_out_ and *A*_in_ respectively denote the surface areas of the outer and inner cortical surfaces.

Across the whole cortex and from the outer to the inner surfaces, we systematically sampled MPI values layer-wise for each of the 64,984 vertices of the HCP template 32 k_LR surface space, which we then averaged within each of the 400 Schaefer parcels, per layer. Following a previously described protocol [33], we constructed subject level 400x400 matrices using pairwise Pearson partial correlation on the MPI profiles of cortical parcels (i.e., correlating the MPI values across 12 layers between parcels), controlling for overall mean cortical MPI, followed by log transformation. We then used these matrices to compute MPC gradients –here directly at the whole brain level instead of independently within hemispheres– by following the same procedure and using the same toolbox and parameters as for computing the FC gradients [30, 31], as described above and previously done [33–35]. We also selected the principal gradient of MPC explaining the most variance in the data, which we labeled the MPC axis and used to represent microstructural organization in subsequent analyses.

### Measures of brain size

In our analyses we included different measures of brain size typically used in the literature, including intracranial volume (ICV), total brain volume (TBV), and total surface area (SA). For ICV, we used the FreeSurfer output measure *IntraCranialVol*, which is an estimate of ICV based on the Talairach transform. We computed our own measure of TBV by summing the volumes of the following FreeSurfer output measures: *TotCort_GM_Vol*, *Tot_WM_Vol*, *3rdVent_Vol*, *L/R_ThalamusProper_Vol*, *L/R_Caudate_Vol*, *L/R_Putamen_Vol*, *L/R_Pallidum_Vol*, *L/R_Hippo_Vol, L/R_Amygdala_Vol*, *L/R_AccumbensArea_Vol*, *L/R_ChoroidPlexus_Vol*, *L/R_LatVent_Vol*, *L/R_InfLatVent_Vol*. We chose to include volumes that are anatomically located within the cortical sheath, which we considered relevant given our study’s focus on cortical functional organization (thus excluding the volumes of subcortical structures). We computed total SA by using the FreeSurfer mri_surf2surf tool to resample cortical white matter surface for each subject.

### Geodesic distances of connectivity profiles

Geodesic distances, representing the shortest distance between two vertices along the folded cortical mantle’s curvature, were computed using the Micapipe toolbox [72], and following the previously described protocol [73]. In short, geodesic distance matrices were computed for each subject along their native cortical midsurface. The first step consisted in defining a centroid vertex for each cortical parcel, identified as the vertex having the shortest summed Euclidean distance from all other vertices within the parcel. Then, Dijkstra’s algorithm [74] was used to compute geodesic distances between the centroid vertices and all other vertices on the on the native midusrface mesh. The vertex-wise geodesic distance values were then averaged within each parcel to form the geodesic distance matrices. From these individual matrices, we finally averaged – parcel-wise– the geodesic distance values of each seed parcel’s top 10% maximally functionally connected regions per hemisphere, thus obtaining for each subject the mean geodesic distance of functional connectivity profiles by region.

### Statistical Analysis

Given that the HCP sample includes different levels of kinship, we used linear mixed effects models (LMMs) to account for sibling status (MZ, DZ, non-twin siblings) and family relatedness. In fact, all LMMs mentioned in this work consistently included sex, age, and total SA as covariates (unless otherwise mentioned), and controlled for random nested effects of family relatedness and sibling status. In addition, effects on cortical data obtained via LMMs underwent false discovery rate (FDR) correction (*q* < 0.05), thus correcting for multiple comparisons across the 400 Schaefer parcels. Throughout this work, we also tested for associations in brain-wide patterns displayed in the form of cortical maps, for which we used Spearman-rank correlation followed by spin-permutation tests to control for spatial autocorrelation [75].

After computing the S-A axis of functional brain organization, we tested for sex differences in the S-A axis loadings with an LMM. Then, we investigated which measure of brain size (out of ICV, TBV, and total SA) had the largest effect on the S-A axis parcel loadings using separate LMMs (respectively only including ICV, TBV, or total SA as a covariate, in addition to sex, age and the random nested effect of family relatedness and sibling status). The reason underlying our decision to systematically include total SA as a covariate in all our LMMs (as the measure of brain size) is that it showed the most widespread effects on the S-A axis loadings out of the three tested measures. Then, we investigated associations between the S-A axis and cortical morphometry, namely the MPC axis and the mean geodesic distance of connectivity profiles, using both LMMs and Spearman-rank correlations of cortical maps.

To probe whether sex differences in cortical morphometry may explain sex differences in the S-A axis, we tested whether sex differences in the S-A axis loadings were moderated by total SA by modelling an additional interaction term of sex by total SA on the S-A axis loadings within the original LMM. We also tested for sex differences in the MPC axis and in the mean geodesic distance of connectivity profiles, and conducted Spearman-rank correlations of cortical *t*-maps for the sex contrast in the S-A axis and in the morphometric measures. Finally, we conducted sensitivity analyses to test for sex effects on the S-A axis yielded by an LMM including all morphometric measures as covariates (i.e., including the MPC axis and the mean geodesic distance of connectivity profiles, in addition to total SA), as well as an LMM not including any morphometric measures as covariates (i.e., also excluding total SA). We then tested the similarity of both these sex effects with the original sex effects on the S-A axis with a Spearman-rank correlation of the cortical *t*-maps.

In order to probe the potential intrinsic functional underpinnings of sex differences in the S-A axis, we tested for sex differences in FC strength (also with an LMM), as well as sex differences in FC profiles, i.e., the presence of sex differences in the top 10% of maximally functionally connected regions used to compute the S-A axis. To this end, we built 400x400 binary matrices at the subject level –based on the subjects’ individual FC matrix z-values– in which we marked in a seed-wise fashion (along the matrix rows) whether the given parcel (along the matrix column) belongs to the given seed’s 10% maximally functionally connected regions, where 1 = parcel belongs to the seed’s top 10% maximally functionally connected regions and 0 = parcel does not belong to the seed’s top 10% of maximally functionally connected regions. We then summed the binary matrices separately within sexes in order to fill 160000 contingency matrices –one for each cell (i.e., functional connection) of the 400x400 FC matrix– as follows:

|  | Parcel belongs to the seed's top 10% of maximally functionally connected regions | Parcel does <u>not</u> belong to the seed's top 10% of maximally functionally connected regions |
| --- | --- | --- |
| Males | $C_m$ | $NC_m$ |
| Females | $C_f$ | $NC_f$ |

where *Cm* and *Cf* respectively denote the number of males and females for which the given parcel (corresponding to the matrix column) constitutes the given seed’s (corresponding to the matrix row) top 10% of maximally functionally connected regions, and where *NCm and NCf* respectively denote the number of males and females for which the given parcel does not constitute the given seed’s top 10% of maximally functionally connected regions.

We then conducted the Chi-square (χ^2^) test of independence (degrees of freedom = 1) on each contingency table to test for sex differences in the odds of each parcel of belonging to the top 10% of maximally functionally connected regions of each seed region. Given the large number of tests conducted here (400x400=160000), we controlled for multiple comparisons using FDR correction. We quantified the size of these sex effects with the odds ratio (OR), calculated as defined in Eq. 2.:

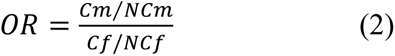

where OR > 1 indicates greater male odds –and OR < 1 indicates greater female odds– of a given region of belonging to a given seed’s top 10% of maximally functionally connected regions.

We also tested for sex differences in netwoexplain sex differences in srk topology, i.e., how nodes are physically organized in networks and how networks are physically organized along the S-A axis. For this, we computed two measures of network dispersion: between-network and within-network dispersion. Between-network dispersion is defined as the Euclidean distance between a pair of network centroids, where a higher value indicates that networks are more segregated from one another along the S-A. Within-network dispersion is defined as the sum squared Euclidean distance of network nodes (i.e., parcel loadings) to the network centroid, where a higher value indicates wider distribution and segregation of a given network’s nodes along the S-A axis. At the individual level, we thus computed between-network dispersion between all networks in a pairwise fashion (21 pairs), and within-network dispersion for all 7 networks, by defining network centroids as the median of the S-A axis loadings of all parcels belonging to a given network, following a previously described protocol [36]. Then, we computed sex differences in each of the 21 between-network dispersion metrics and 7 within-network dispersion metrics using LMMs. For each model, we computed a null distribution of *t*-values for sex differences using 1000 spherical rotations of the Schaefer parcellation scheme in order to shuffle the network labels [75], against which we computed our *p*-value to determine statistical significance. We then assed *p*_spin_-values against Bonferroni-corrected two-tailed α-levels of 0.001 (0.025/21) and 0.004 (0.025/7) for between-network and within-network dispersion sex contrasts respectively.

## Author contributions

Conceptualization: BS and SLV. Main analysis and visualization: BS. Input on analysis: MDH, GB, and SLV. Writing—original draft: BS. Writing—review and editing: BS, MDH, LW, GB, JS, SW, SBE, SLV. Supervision: SLV.

## Data and materials availability

All data needed to evaluate the conclusions in the paper are present in the paper and the Supplementary Materials. We obtained human data from the open-access Human Connectome Project HCP S1200 young adult sample. Data are available upon request at http://www.humanconnectome.org/. Analyses were conducted in Python and R: The code used in this manuscript is available at https://github.com/biancaserio/sex_diff_gradients. The code and tutorials for functional gradient decomposition and to generate geodesic distances can further be found at https://brainspace.readthedocs.io/en/latest/index.html and https://micapipe.readthedocs.io/en/latest/ respectively.

## SUPPLEMENTARY MATERIALS

Supplementary results and methods can be found in the Supplementary Materials.

