## Supplementary Materials for "Sex differences in intrinsic functional cortical organization reflect differences in network topology rather than cortical morphometry"

**This PDF file includes:**

Supplementary Results
Figs. S1 to S4
Tables S1 to S2

Supplementary Methods
Text
Figure S5

### Supplementary Results

#### Figures

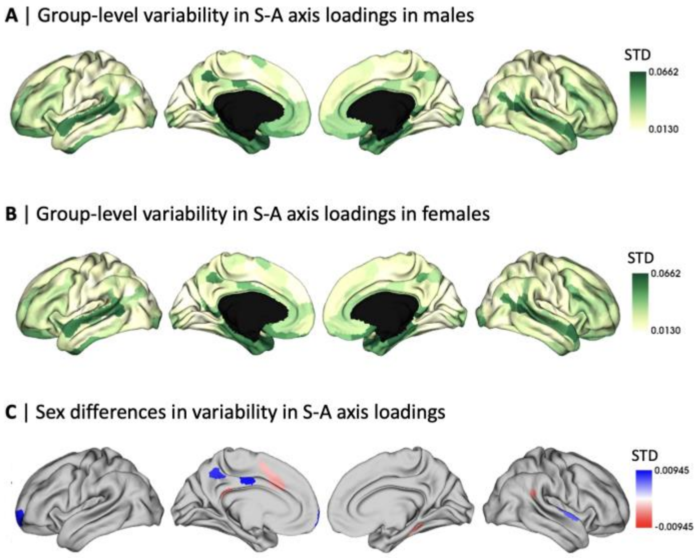


**Figure S1. Variability in the sensory-association (S-A) axis of functional cortical organization.** Group-level variability (quantified by standard deviation; STD) in S-A axis loadings in **A** | males and **B** | females; **C |** Thresholded (false discovery rated (FDR)-corrected, $q < .05$) map of sex differences in variability (STD) as determined by Levene's test for equality of variances, where blue represents higher male variability and red represents higher female variability.

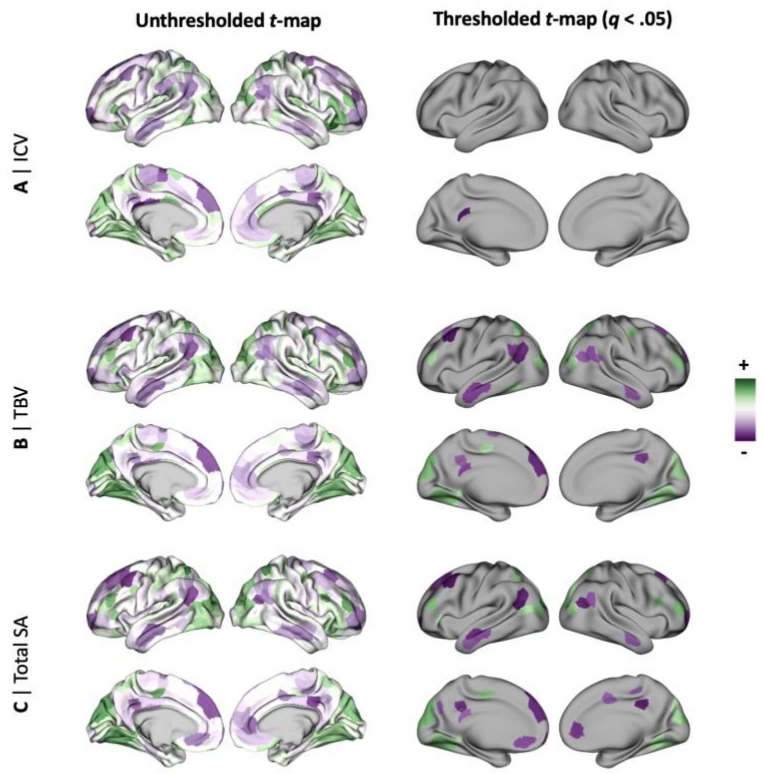


**Figure S2. Effects of brain size on the sensory-association (S-A) axis of functional cortical organization.** Unthresholded and thresholded (false discovery rated (FDR)-corrected, $q < .05$) $t$-maps of linear mixed effects model results showing the effects of different measures of brain size, namely **A |** intracranial volume (ICV), **B |** total brain volume (TBV), and **C |** total surface area (SA), on S-A axis loadings. Total SA yielded the largest number of significant parcels following FDR correction (65), followed by TBV (59), and finally ICV (2).

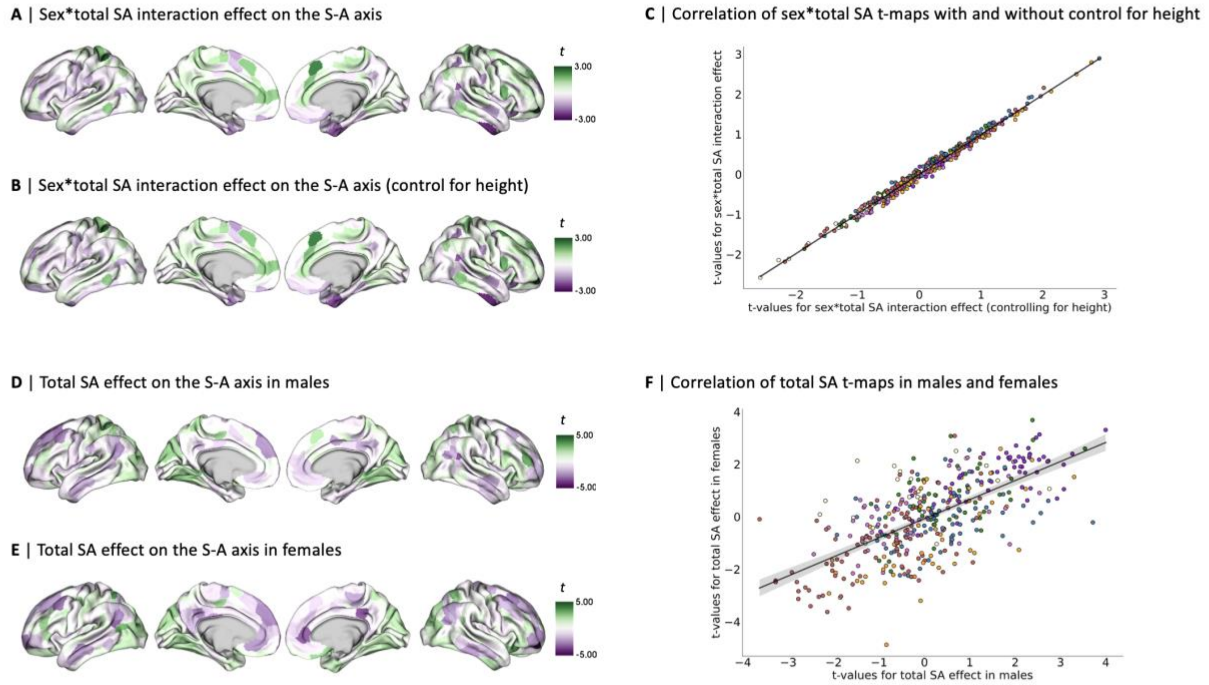


**Figure S3. Effect of total surface area (SA) on the sensory-association (S-A) axis of functional cortical organization and its relationship to sex. A |** Unthresholded *t*-map of linear mixed effect model (LMM) testing for sex by total SA interaction effect on S-A axis loadings (original result, shown in Figure 3B); **B |** Unthresholded *t*-map of LMM testing for sex by total SA interaction effect on S-A axis loadings when including height as a covariate in the LMM; **C |** Scatterplot displaying the spatial correlation between patterns of sex*total SA interaction effects on S-A axis loadings with (x-axis) and without (y-axis) controlling for height in the LMM (color-coded by yeo network), $r$ = 0.99, $p_{spin} < .001$; **D |** Unthresholded *t*-map of LMM testing for sex effect on S-A axis loadings in males; **E |** Unthresholded *t*-map of LMM testing for sex effect on S-A axis loadings in females; **F |** Scatterplot displaying the spatial correlation between patterns of total SA effects on S-A axis loadings in males (x-axis) and females (y-axis) (color-coded by yeo network), $r$ = 0.65, $p_{spin}$ = .001.

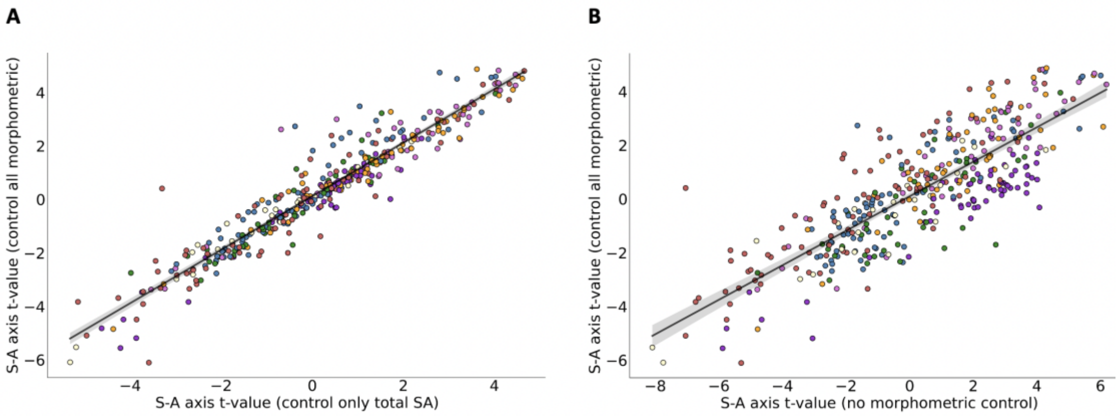


**Figure S4. Similarity of sex differences in the sensory-association (S-A) axis with and without controlling for morphometric measures.** Scatterplots displaying the spatial correlation between patterns of sex effects on S-A axis loadings with (y-axis) and without (x-axis) inclusion of morphometric measures as covariates in the LMMs (color-coded by yeo network), where **A** | LMM not including morphometric measures (x-axis) still includes total SA as a covariate (model used to yield main sex difference results, Fig. 1B), $r$ = 0.95, $p_{spin}$ < .001. **B** | LMM not including morphometric measures (x-axis) does not include any morphometric measure as covariates, $r$ = 0.81, $p_{spin}$ < .001.

#### Tables

| | Male | | Female | | $t$ |
|---|---|---|---|---|---|
| | **Mean** | **SD** | **Mean** | **SD** | |
| **ICV ($cm^3$)** | 1'710.35 | 140.91 | 1'474.63 | 149.86 | 27.45** |
| **TBV ($cm^3$)** | 1'099.51 | 92.35 | 958.10 | 81.25 | 29.98** |
| **Total SA ($cm^2$)** | 1'947.35 | 1'542.24 | 17'150.67 | 1'469.42 | 27.43** |
| **Height (in)** | 70.47 | 2.91 | 64.91 | 2.65 | 36.27** |
| **Weight (lbs)** | 189.93 | 34.75 | 156.19 | 35.82 | 15.35** |
| **BMI** | 26.84 | 4.33 | 26.04 | 5.66 | 2.50* |

**Table S1. Sex differences in brain size and anthropometric measurements.** Results are yielded by the linear mixed effects model: brain size/anthropometric measurement ~ 1 + sex + age + (1 | family relatedness / sibling status). * indicates $p < 0.05$, ** indicates $p < .001$. SD, standard deviation; ICV, intracranial volume; TBV, total brain volume; SA, surface area; BMI, body mass index.

| | $t$ | $p$ | $p_{spin}$ |
|---|---|---|---|
| **WN dispersion (network)** | | | |
| Visual | -3.033 | 0.002 | 0.011 |
| Somatomotor | 3.103 | 0.002 | 0.004 |
| Dorsal attention | -0.715 | 0.475 | 0.237 |
| Ventral attention | 2.388 | 0.017 | 0.020 |
| Limbic | 1.963 | 0.050 | 0.111 |
| Fronto parietal | 1.431 | 0.152 | 0.076 |
| DMN | 2.412 | 0.016 | 0.001* |
| **BN dispersion (pairwise networks)** | | | |
| Visual - somatomotor | 1.394 | 0.163 | 0.322 |
| Visual - dorsal attention | 0.678 | 0.498 | 0.421 |
| Visual - ventral attention | -1.925 | 0.054 | 0.249 |
| Visual - limbic | 2.597 | 0.009 | 0.19 |
| Visual - fronto parietal | -1.505 | 0.132 | 0.298 |
| Visual - DMN | 1.070 | 0.285 | 0.377 |
| Somatomotor - dorsal attention | -0.666 | 0.505 | 0.372 |
| Somatomotor - ventral attention | -2.990 | 0.003 | 0.061 |
| Somatomotor - limbic | 2.263 | 0.024 | 0.258 |
| Somatomotor - fronto parietal | -2.536 | 0.011 | 0.113 |
| Somatomotor - DMN | 0.131 | 0.896 | 0.494 |
| Dorsal attention - ventral attention | -3.111 | 0.002 | 0.100 |
| Dorsal attention - limbic | 2.191 | 0.028 | 0.255 |
| Dorsal attention - fronto parietal | -2.229 | 0.026 | 0.202 |
| Dorsal attention - DMN | 0.593 | 0.553 | 0.432 |
| Ventral attention - limbic | 3.331 | 0.001 | 0.101 |
| Ventral attention - fronto parietal | 0.385 | 0.700 | 0.413 |
| Ventral attention - DMN | 2.744 | 0.006 | 0.100 |
| Limbic - fronto parietal | -3.238 | 0.001 | 0.118 |
| Limbic - DMN | -2.176 | 0.030 | 0.218 |
| Fronto parietal - DMN | 2.832 | 0.004 | 0.090 |

**Table S2. Sex differences in within-network (WN) and between-network (BN) dispersion.** Results are yielded by the linear mixed effects model: dispersion ~ 1 + sex + age + total surface area + (1 | family relatedness / sibling status). * indicates statistical significance at Bonferroni-corrected thresholds of 0.0036 for WN dispersion (7) comparisons, and 0.001 for BN dispersion (21) comparisons. DMN, default mode network.

#### Supplementary Methods

Concerns have been recently expressed regarding the reliability of the T1w/T2w ratio to quantify microstructural profile intensity (MPI). In fact, MPI is subject to a B1 field bias that has been shown to correlate with demographic and compositional variables such as age, sex, and body mass index (BMI), potentially leading to spurious results when statistical comparing MPI across individuals and groups [77]. However, we did not expect that bias in MPI values would persist in our MPC axis, given that it is an inherently relative measure of intra-individual variation, which is further computed by regressing out mean cortical MPI. Indeed, we show in Supplementary Fig. S5 – via correlations between MPI/MPC axis and BMI/ICV– that biases observed when plotting MPI as a function of BMI and ICV (i.e., the distributions of correlation coefficients r(MPI, BMI) and r(MPI, ICV) across sexes are skewed) are not observed when plotting MPC as a function of BMI and ICV (i.e., the distributions of correlation coefficients r(MPC, BMI) and r(MPC, ICV) are normal and do not vary as a function of sex). As such, we confirmed the suitability of MPI, yielded by the T1w/T2w ratio, to compute the MPC axis for further analyses assessing its variation between sexes without introducing bias.

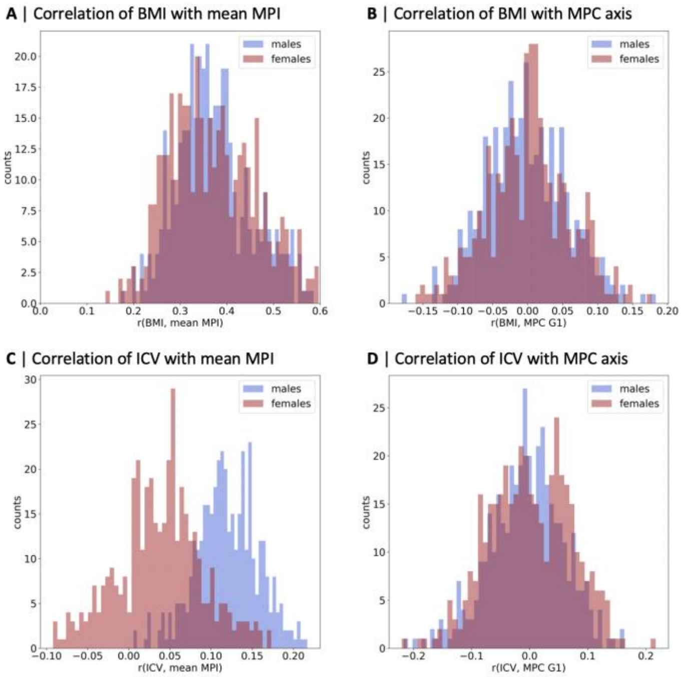


**Figure S5. Check for field bias in T1w/T2w raw mean microstructure profile intensity (MPI) and derived microstructure profile covariance (MPC) axis.** Histograms of correlation coefficients (color-coded by sex) for correlations between: **A** | Body mass index (BMI) and mean MPI (colored by sex); **B** | BMI and mean MPC axis loadings; **C** | Intracranial volume (ICV) and mean MPI (colored by sex); **D** | ICV and mean MPC axis loadings.
